# Segmental duplication-mediated rearrangements alter the landscape of mouse genomes

**DOI:** 10.1101/2025.07.18.665526

**Authors:** Eden R. Francoeur, Peter A. Audano, Ardian Ferraj, Parithi Balachandran, Christine R. Beck

## Abstract

Segmental duplications (SDs) are dynamic regions of mammalian genomes that drive structural variation and phenotypic diversity. However, genome-wide characterization of the SD landscape and the rearrangements mediated by SD paralogs in mouse genomes has been limited by the near-exclusive reliance on short-read sequencing technologies. Here, we integrate long-read genome assemblies, optical mapping, and k-mer based copy number analysis across eight genetically diverse inbred mouse strains to identify and characterize SD-mediated rearrangements. We identify 223 rearrangements affecting over 14 Mb of sequence and reveal substantial variation in gene content. These rearrangements affect loci involved in immunity, sensory perception, and gene regulation, including variation in the amylase gene cluster (*Amy2a*) and KRAB-zinc finger genes. We observed that SD flanks in mouse genomes are significantly enriched for young LINE-1 transposable elements, suggesting a potential role for transposons in promoting recombination and generating SDs within mice. Our findings highlight the contribution of SDs to genome structure and intra-species variation, and provide a resource for identifying regions prone to rearrangement in a critical model organism for biomedical research.

## INTRODUCTION

Segmental duplications (SDs) are among the most rapidly evolving regions of mammalian genomes and can generate significant variation within species [1–3]. These duplications are defined as large (≥1 kb) and highly homologous (≥90%) DNA sequences excluding transposable elements. SDs constitute over 5% of both human and mouse genomes and frequently contain genes [3–5]. Due to the homology and length of SDs, they can undergo ectopic rearrangement resulting in large structural variants (SVs; ≥50 bp) such as deletions, duplications, and inversions depending on the sequence orientation of the SD paralogs with respect to each other. These ectopic rearrangements typically occur through non-allelic homologous recombination (NAHR), a mechanism that is thought to require 200 base pairs of perfect homology, although other mechanisms can also generate rearrangements flanked by SDs [6–12].

SD-mediated rearrangements play an important evolutionary role by contributing to genetic diversity and phenotypic variation. Changes in gene dosage can serve as targets for adaptive evolution, as seen by copy number polymorphisms of amylase genes within mammalian genomes [13–15]. Some SD-mediated rearrangements are responsible for human diseases, including Prader-Willi syndrome [16], Charcot-Marie-Tooth disease (through duplication of *PMP22*), and hereditary neuropathy with pressure palsies (through deletion of *PMP22*) [17]. Other SD-mediated rearrangements in the human genome are implicated in variation in sensory perception, reproduction, and behavior, which can also lead to fitness differences [18–22].

Mice are an important model organism used to study human disease, and understanding genetic variation between strains is critical for interpreting results, experimental design, and generating appropriate comparisons. Recent efforts have revealed the role of SD-mediated rearrangements in shaping genetic diversity in mouse genomes through manual assembly and curation to investigate copy number variation [23]. These SD-mediated rearrangements result in strain-specific haplotypes containing genetic variation in loci implicated with sensory, behavior, and immunity related differences, including variability within the defensin gene family, which is associated with differences in infection susceptibility across mouse strains [4, 23]. SD-mediated variation may also contribute to male sterility when PWD/PhJ mice are crossed with C57BL/6J due to an SD-mediated deletion of the *Hstx2* region in PWD/PhJ [24]. While these site-specific analyses have depicted the importance of SD-mediated variation in mouse strains, a genome-wide examination has not yet been conducted because of the limitations of existing references and short-read sequencing. With advances in long-read sequencing technology and whole-genome assembly, we can now accurately examine repetitive sequences within genomes and retrieve almost 3× more SVs compared to short-reads [25–28].

By combining long-read whole-genome assemblies, optical mapping, and short-read sequencing, we identify SD variation across the genomes of eight inbred mouse strains important for biomedical research. We investigate how SDs drive SV formation within strains from the *Mus musculus* species that span ∼1/2 a million years of evolution. Our analyses reveal SD characteristics leading to recombination, identify regions prone to future rearrangements, and show how SDs act as drivers of genomic and transcriptomic evolution within a species. In total, more than 14 Mb of genomic differences between strains can be attributed to SD-mediated rearrangements. As SDs underlie both adaptive traits and pathogenic rearrangements in humans, a deeper understanding of their behavior in mice offers insights into mutational mechanisms and evolutionary forces that influence diversity, genome stability, and disease susceptibility. Because SD-mediated rearrangements directly impact neurodevelopmental, immunological, cardiovascular, and other disorders and phenotypes, identifying existing rearrangements and predicting sites prone to *de novo* rearrangements is important for biomedical research utilizing mouse models [29–32].

## RESULTS

### The segmental duplication annotation and landscape of GRCm39

We previously generated whole-genome assemblies and a high-quality SV callset for multiple diverse mouse genomes [25]. These assemblies have enabled the investigation of regions of the genome that were previously inaccessible. To use these assemblies to examine SD-mediated variation across mouse strains, we first had to establish the SD landscape of the GRCm39 mouse reference genome. In order to do so, we compared the SD annotation results from SEDEF and BISER with WGAC SD annotations in GRCm38, which was previously considered as the gold standard (Methods). SEDEF did not identify ∼10% of SD sequence identified with WGAC, compared to BISER, which missed ∼25% of WGAC SD sequence and produced many more short annotations (Supp 1A-B). We therefore annotated SDs in GRCm39 (mm39) using SEDEF, including only SDs ≥ 90% identity to their paralog and ≥ 1 kb in length (Methods) [5, 33]. Our analysis identified 198,888 SDs in the GRCm39 genome. Of these, we find 54,730 on the same chromosome (intrachromosomal), with 32,744 in direct orientation and 21,986 in an inverted orientation. Intrachromosomal SD paralogs also tend to exhibit significantly higher identities and lengths, averaging 25-30 kb longer than interchromosomal SDs (Fig 1A-B). The length of SDs varies from 1 kb to 129 kb (median: 2,783 bp) for interchromosomal SDs and from 1 kb to over 1 Mb (median: 13,616 bp) for intrachromosomal SDs (Supp 1C). In this work, we characterize intrachromosomal SD paralogs and their effects on intrachromosomal rearrangements, which also rearrange at a higher rate [34].

**Figure 1.**
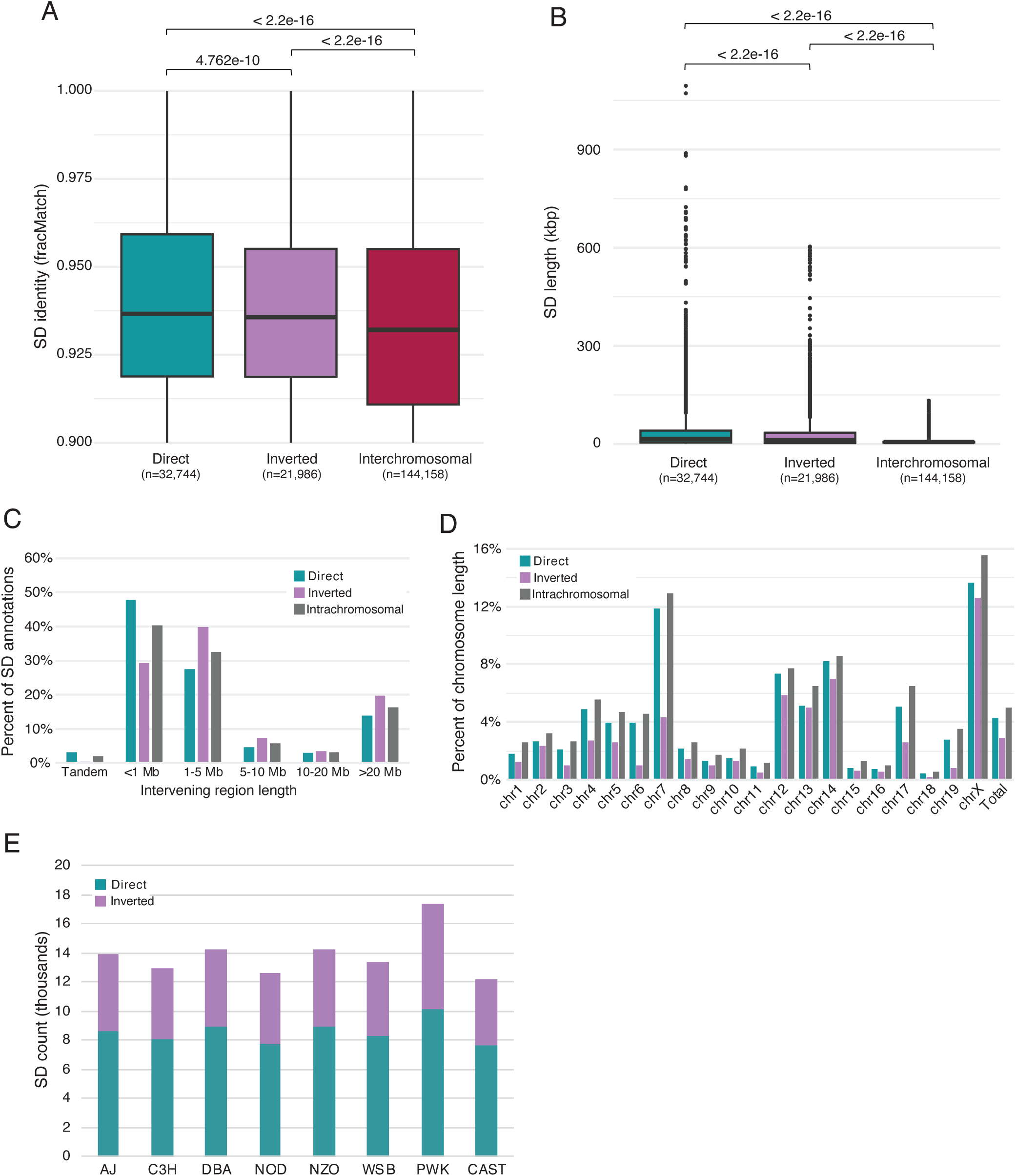
**SD annotation and landscape in GRCm39 (**A) Percent identity of SDs identified by SEDEF in GRCm39. SDs are grouped into three categories: those in direct orientation to their paralog (teal), those in inverted orientation with their paralog (purple), and interchromosomal (red) (Mann-Whitney U test). (B) Length of GRCm39 SDs. SDs are grouped the same as in (A) (Mann-Whitney U test). (C) Distribution of the distance between (intervening region) direct and inverted SD paralogs identified in GRCm39. (D) Percent of chromosome length composed of intrachromosomal SDs in GRCm39. Direct and inverted SDs may overlap, gray bars show the SD content of each chromosome when direct and inverted SDs are merged. (E) Intrachromosomal SDs annotated by SEDEF in eight mouse strains: A/J (AJ), C3H/HeJ (C3H), DBA/2J (DBA), NOD/ShiLtJ (NOD), NZO/HILtJ (NZO), WSB/EiJ (WSB), PWK/PhJ (PWK), and CAST/EiJ (CAST).

Further examination of GRCm39 intrachromosomal SD annotations reveals that direct SDs tend to have shorter intervening sequence lengths compared to inverted SDs. Approximately 49% of direct and 70% of inverted SD paralogs have ≥ 1 Mb of sequence between paralogs, indicating potential differences in the mechanism(s) of their formation or their tolerance in the genome (Fig 1C). This difference could also lead to functional impacts and constraints on SV formation and stability depending on their orientation. This result is consistent when examining GRCm39 SDs 10 kb in length or greater (Supp 1D).

The distribution of SDs across the mouse genome is not uniform, with over 5% of GRCm39 composed of intrachromosomal SD paralogs. Chromosome 7 is enriched in directly oriented paralogs that make up ∼12% of the chromosome (Fig 1D). Similarly, ∼15.5% of the mouse X

Chromosome is spanned by SDs, which is a larger portion than any autosome and has an abundance of inverted SDs (Fig 1D), potentially indicating an increased predisposition to inversions on this chromosome [35, 36]. This differs from the human X chromosome which contains a moderate amount of SDs in comparison [3].

We next annotated SDs in unphased, haploid long-read genome assemblies for eight strains of inbred mice [25], including six founders of the Collaborative Cross, a panel of recombinant inbred mice derived from multiple parental strains (A/J, CAST/EiJ, NOD/ShiLtJ, NZO/HILtJ, PWK/PhJ, and WSB/EiJ) [37] and two additional strains that are commonly used models (C3H/HeJ and DBA/2J). We produced SD annotations using the same methods as GRCm39 (Methods). On average *Mus musculus domesticus* strains have 13,576 (45.5 Mb) intrachromosomal SDs, while PWK/PhJ (*M. m. musculus)* has 17,375 (37.3 Mb), and CAST/EiJ (*M. m. castaneus*) has 12,249 (41.3 Mb) (Fig 1E, Supp 1E). Although the number of SDs across our eight assemblies are comparable, we identified fewer SDs in the long-read assemblies compared to the GRCm39 reference. This discrepancy likely reflects the lower contiguity of the assemblies in comparison to the reference (Methods). While PWK/PhJ has the largest number of SDs, its overall SD content is similar to other strains (Supp 1F). Our SD annotations provide a comprehensive map of the duplication landscape across multiple mouse genomes.

### Transposable elements are prevalent at SD boundaries

NAHR between repetitive DNA sequences, such as transposable elements (TEs) and simple repeats (SR), has previously been implicated in the formation of large duplications or SDs [38, 39]. To understand the mechanisms underlying SD formation in mouse genomes, we analyzed the repetitive sequence classes at the boundaries of mouse SDs and compared our findings to human genomes (Methods).

While human genomes are enriched in *Alu* TEs at the flanks of SDs [38], when examining the direct intersection (within 10 bp of the SD coordinates) of SD boundaries from GRCm39 with RepeatMasker (Methods), we found that the SINE B2 and B4 mouse TEs, which are closely related to *Alu* elements, are significantly depleted at the boundaries of SDs (*P*-value ≤ 0.001, permutation test) (Fig 2A).

**Figure 2.**
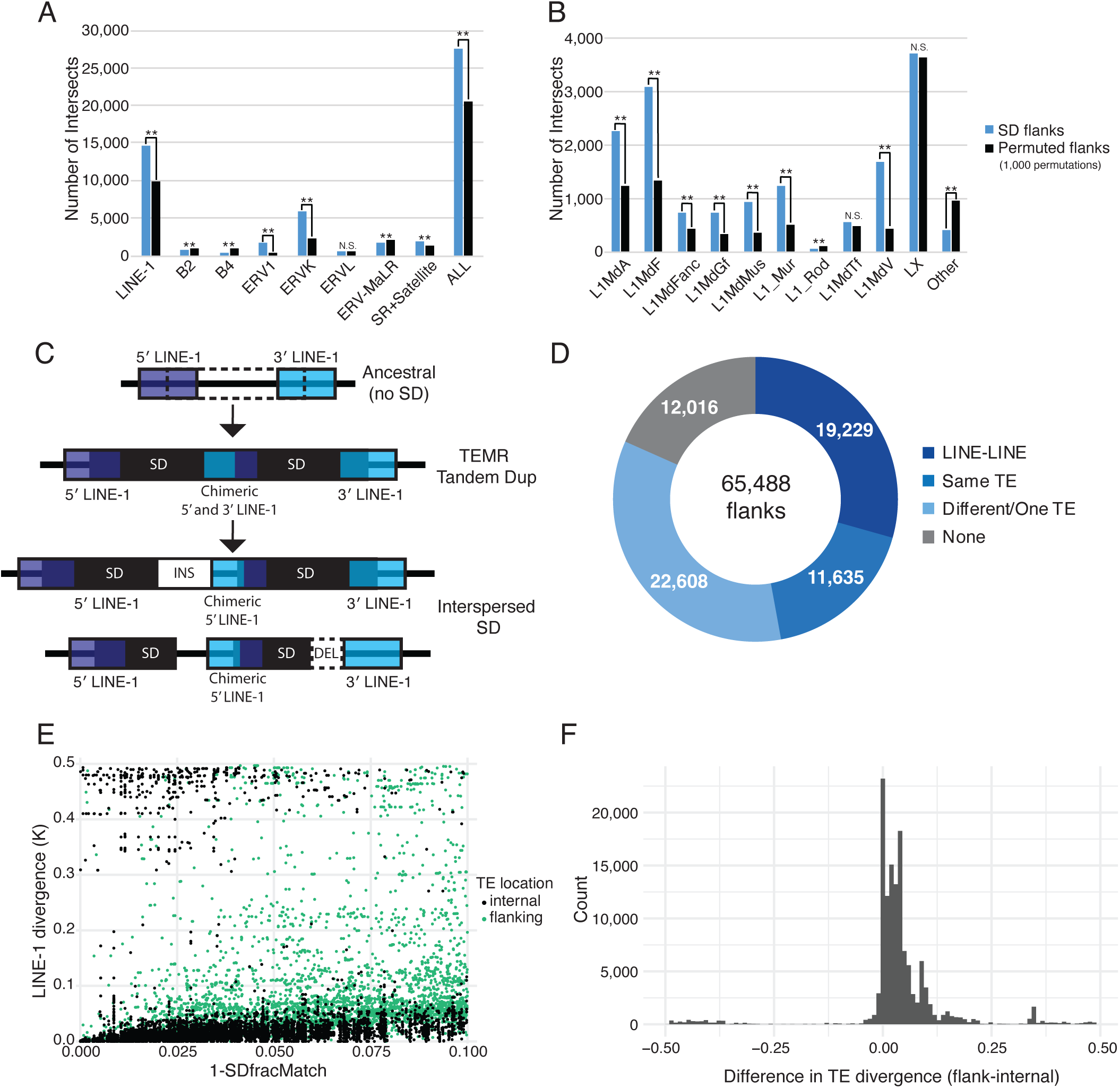
Origin of SDs in GRCm39. (A) Number of intersects of SD flanks with TE types compared with 1,000 permutations of flanking regions (** = p ≤ 0.001, N.S = not significant). (B) Number of intersects of SD flanks with LINE-1 subtypes compared with 1,000 permutations of flanking regions (** = p ≤ 0.001, N.S. = not significant). (C) Ectopic repair between transposable elements may seed SD formation. Shown is a cartoon depicting how TEs can potentially underlie both tandem and interspersed SD formation. For tandem duplications, the 5′ and 3′ LINE-1s flanking both SD copies remain the same as the ancestral state, and there is a chimeric LINE-1 in the middle of the two paralogs. At the bottom, we show that insertions or deletions occurring within SDs can cause interspersed duplications where one SD paralog may not have both 5′ and 3′ TEs. (D) Count of SD flanks intersecting the same or different TE types for direct SDs. 5′ flanks were compared to 5′ flanks (and 3′ to 3′) between direct SD paralogs and 5′ to 3′ (and 3′ to 5′) comparisons for inverted SD paralogs. LINE-LINE: LINE TEs at both flanks, Same: same TE type at flanks (excluding LINE-LINE), Different/One TE: Different TE types at flanks or TE at only one flank, None: no TEs at either flank. (E) The sequence divergence of LINE-1s at flanks of direct SD paralogs (5′ to 5′ and 3′ to 3′ comparisons) and TEs internal to direct SDs compared to SD divergence by Kimura’s two-parameter distance model. (F) Difference in the sequence divergence of flanking vs internal LINE-1 elements.

Our analysis identified a significant enrichment of LINE-1, ERV1, and ERVK TEs at the flanks of SDs in mouse genomes, with LINE-1s occurring most often (*P*-value ≤ 0.001, permutation test) (Fig 2A). Simple and satellite repeats were also moderately enriched. We next compared the enrichment of LINE-1 elements at the flanks of direct and inverted SD paralogs. Given that direct SDs are often separated by shorter distances than inverted SDs, we hypothesized that they may be more likely to arise through NAHR mediated by LINE-1 elements. Supporting this, we observe significant LINE-1 enrichment at the flanks of both orientations, with a stronger enrichment at direct SDs (Z-score: 8.3882) compared to inverted SDs (Z-score: 5.465) (Supp 2A).

We find that most LINE-1 subfamilies are significantly enriched at SD flanks, with the exception of Lx, L1MdTf, L1_Rod, and a few other unclassified LINE-1 elements that did not fit into the main LINE-1 subfamilies (Fig 2B). While it is the most abundant LINE-1 subtype in GRCm39, Lx was not significantly enriched at SD flanks. In contrast, L1MdA and L1MdF, which were the most enriched at SD flanks, are among the more abundant LINE-1 subfamilies (Supp 2B).

We next hypothesized that the LINE-1 enrichment at SD flanks is influenced by the mean length of elements in the subfamilies, which may increase their capability to mediate rearrangements through NAHR, by providing long stretches of perfect homology. Our analysis confirmed that L1MdTf elements (mean 2,417 bp) are longer than other subfamilies, and L1MdA (mean 1,385 bp), L1MdF (mean 1,108 bp), and L1MdGf (mean 1,633 bp) elements are longer than Lx (mean 502 bp) elements (Supp 2C). However, despite the greater length of L1MdTf elements, this did not result in their enrichment at SD flanks (Fig 2B, Supp 2C), indicating that element length is likely not the sole factor for LINE-1 enrichment at the border of SDs.

In addition to their length and frequency, we next considered whether the age and therefore divergence of LINE-1 elements within a subfamily could also influence their enrichment at SD flanks; in humans, only *Alu*S and *Alu*Y subfamilies are enriched at SD boundaries. While older LINE-1 subfamilies were once young and more identical, and therefore potentially capable of NAHR, their lack of enrichment at current SD flanks may reflect their inability to act as substrates for NAHR or that the SDs they nucleated have accumulated mutations and no longer meet the definition of an SD. The most recently active subtype, L1MdTf (active from 0-1 MY ago), was not significantly enriched at SD flanks, suggesting that its recent activity may not have allowed enough time for it to contribute to SD formation or did not coincide with evolutionary pressures to form SDs. The relatively younger LINE-1 elements, L1MdA, L1MdF, and L1MdGf (0-6 MY ago), have been present in mouse genomes for longer than L1MdTf, and all three of these classes were significantly enriched at SD flanks. Significantly older LINE-1 subtypes, such as L1MdFanc, L1_Mus, and L1MdV were less enriched at SD flanks (6-11 MY ago), which could reflect reduced capacity to mediate recent rearrangements resulting in SDs. Lx elements date back to the split of mice and rats ∼13 million years ago and are both shorter and more diverged [40–42].

### SDs may have arisen by NAHR between homologous TEs

The enrichment of LINE-1 TEs at the boundaries of SDs suggests that homology-driven mechanisms including double-strand break repair and NAHR are responsible for SD formation. Therefore, we propose that for either tandem SDs or interspersed intrachromosomal SDs that ancestral TEs that could predispose the locus to NAHR or other homology directed repair mechanisms [7, 8, 10, 38, 43]. TEs could nucleate tandem and interspersed SDs by homology directed repair followed by divergence of the duplicate locus is depicted (Fig 2C), although other mechanisms involving extrachromosomal DNA have been proposed [38]. To test if paralogous TEs mediated rearrangements resulting in SD formation, we examined direct SD paralogs for matching TEs at the 5′ and 3′ flanks (5′ to 5′ and 3′ to 3′ comparisons) (Methods). While the 5′ and 3′ ends of a majority of SD paralogs intersected TEs, about half of the flanks had mismatching TE types (e.g., SINE at the first paralogs 5′ flank and LINE at the second paralogs 5′ flank) or only had a TE at the flank of one paralog (Fig 2D). SD paralogs with the same TE type at both of the paralog’s flanks were predominantly LINE-1 elements (>60%). These LINE-1 elements have an average length of 2,062 bp, which is significantly larger than the average of all GRCm39 LINE-1 elements (598 bp) and all L1Md elements (1,135 bp). We found that 64% of SD paralogs with LINE-1 elements at both 5′ or both 3′ flanks are L1Md elements (*P*-value < 2.2e-16, Fisher’s Exact Test; odds ratio = 40.7).

To test if SD formation is due to rearrangements between these flanking LINE-1s, we compared the divergence of flanking LINE-1 elements to internal LINE-1 elements (Methods). LINE-1 elements internal to the SD are less diverged in comparison to flanking LINE-1 elements (2.9% internal, 8.7% external, *P*-value <2.2e-16, Wilcoxon rank-sum test) (Fig 2E, Supp 2D). We examined the difference in divergence between flanking and internal TEs (flank-internal) in a pairwise manner (Fig 2F) and find that the flanks of SD paralogs are more diverged than the TEs internal to them (*P*-value <2.2e-16, Wilcoxon signed-rank test). The higher divergence observed in flanking TEs mirrors that of human *Alu* elements flanking SDs vs the divergence *Alu* sequences internal to SDs [38]. Overall, our data provide evidence that paralogous TEs (primarily LINE-1 sequences) underlie the formation of paralogous SD copies in mouse genomes, analogous to *Alu* sequences in human genomes (Fig 2C).

### Genome-wide predictions of SDs susceptible to NAHR in the *Mus musculus* genome

Using our GRCm39 SD annotations, we next predicted regions susceptible to SD-mediated rearrangements across the mouse reference genome to identify regions that may predispose animals to disease or phenotypic differences. Using previously determined parameters associated with human and mouse rearrangements [1, 4, 23] and with predictive capabilities for NAHR hotspot prediction [18, 44], we identified GRCm39 intrachromosomal SD paralogs with ≥95% sequence identity that were greater than 1 kb and had less than 10 Mb of intervening sequence (Methods) (Supp 3A). Our analysis indicates that 29.2% of SD paralogs in direct orientation and 23.7% in inverted orientation are candidates for NAHR. We call the SDs likely to underlie rearrangements “predicted SD paralogs” (Fig 3A).

**Figure 3.**
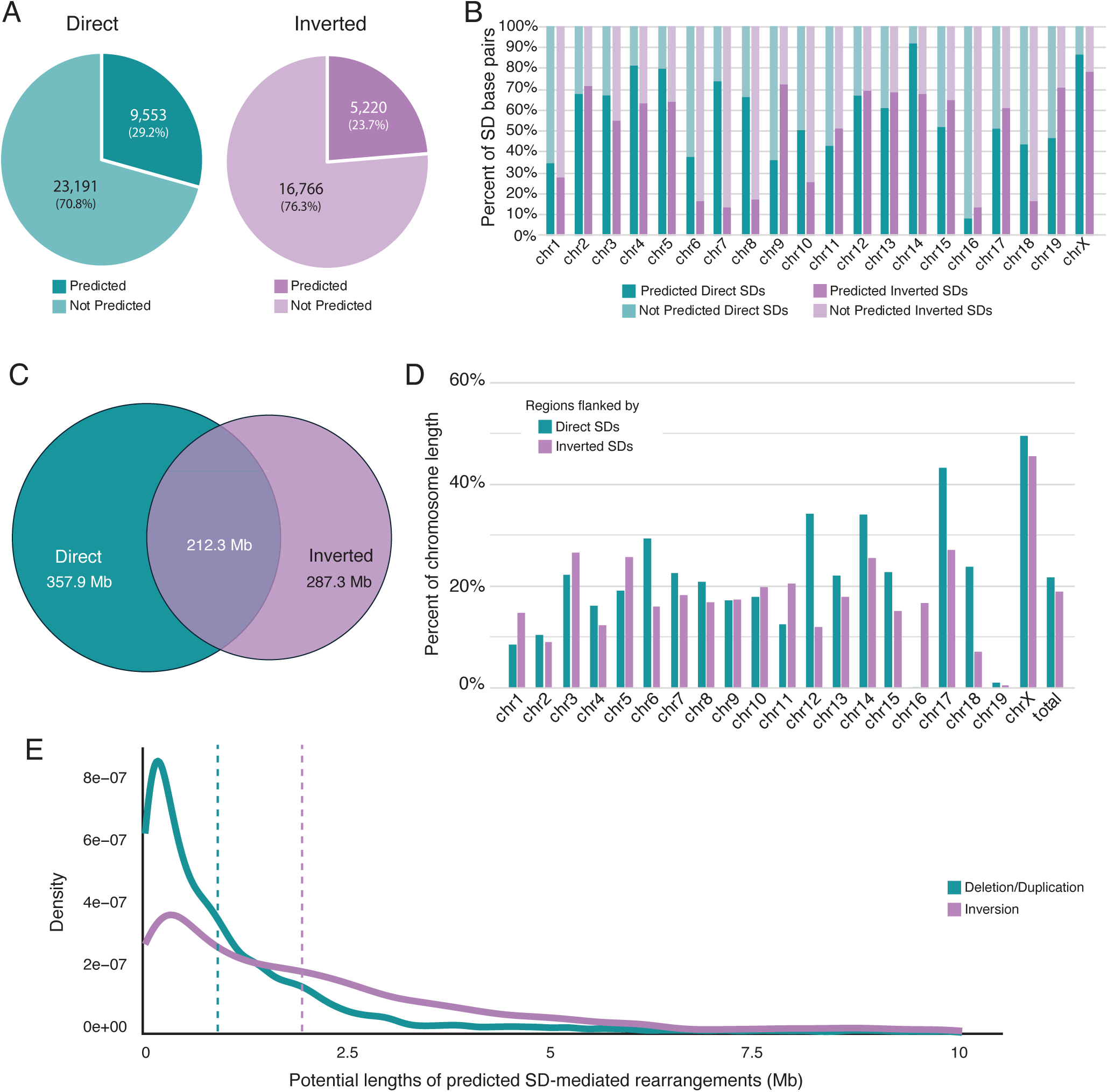
Prediction of SD recombination hotspots in GRCm39. (A) Percent of direct (teal) and inverted (purple) SDs that are predicted to have the potential to undergo rearrangements. (B) Percent of SDs on each chro-mosome that are predicted to have the potential to undergo rearrangement. (C) Regions of the genome prone to deletions and duplications, inversions, or both. (D) Percent of each chromosome that has the potential to undergo a rearrangement (percent of each chromosome flanked by predicted SD paralogs). (E) Distribution of lengths of potential rearrangements mediated by predicted direct (teal) and inverted (purple) SDs (Welch Two Sample t-test: p-value < 2.2e-16). Dotted lines indicate means.

We find that the percentage of predicted SD paralog base pairs relative to the total number of SD base pairs varies across chromosomes. Chromosomes 6, 7, and 8 have less than 20% of inverted SDs matching the prediction criteria, whereas almost all SDs on the X Chromosome have the potential to recombine using our prediction metrics (Fig 3B). We find 570.3 Mb (21.7% of the genome, excluding the Y Chromosome) is flanked by predicted SD paralogs and may be susceptible to NAHR-mediated deletions and duplications. Meanwhile, 499.7 Mb of GRCm39 (19%) is flanked by inverted predicted SD paralogs predisposing them to inversions. In total 857.6 Mb (32.6%) of the mouse reference genome is susceptible to SD-mediated rearrangements by either direct or inverted paralogs and 212.4 Mb (8.1%) is flanked by both types of predicted repeats (Fig 3C). Notably, 49.5% and 45.5% of the X Chromosome is flanked by direct and inverted predicted SD paralogs, respectively (Fig 3D). Although Chromosome 16 contains few predicted SD paralogs (Supp 3B), over 15% of the chromosome has the potential for recombination between inverted SDs, indicating that these predicted SDs are distal to their paralogs. Interestingly, the predicted directly oriented SD paralogs on Chromosome 16 flank relatively small regions, suggesting that they are likely tandemly repeated.

While a majority of the predicted SD paralogs are <2 Mb apart (mean 1.2 Mb), inverted SD paralogs tend to be significantly further apart (mean 2.1 Mb) (*P*-value <2.2e-16, Welch’s two-sample *t*-test) (Fig 3E).

To find loci unlikely to tolerate SD-mediated rearrangements, we next identified regions that might impact mouse viability or fitness. We find 5,558 protein coding genes within the intervening regions of directly oriented predicted SD paralogs and 731 genes that lie within the inverted predicted SDs that could be disrupted by an inversion (Methods). These include transmembrane signal receptors, including olfactory receptor genes, transcriptional regulators, such as KRAB zinc-finger protein genes, and immune related genes (Supp 3C-D). We did not consider genes that lie within the intervening region of predicted inverted paralogs even though these may be affected by the disruption of spacing of their local genomic context or *cis-*regulatory elements.

We identified 914 (16.5%) genes in the MGI database of mammalian phenotypes [45, 46] that reside between directly oriented predicted SD paralogs (Methods) where a gene deletion may be lethal (Supp 3C). Genes identified within these SD bounded loci include *Eif2b3*, which encodes for a subunit for the translation initiation factor Eif2b, and Braf, which encodes a protein involved in cell growth and division [47–49]. Moreover, 45 of the 731 (6.1%) genes that lie within oppositely oriented predicted SDs may also result in lethality (Supp 3D). While these loci may be susceptible to NAHR, rearrangements involving these genes are less likely to be segregating in mouse strains, even in a laboratory setting.

### SD-mediated rearrangements diversify inbred mice

To accurately identify SD-mediated rearrangements across diverse mice, we utilized three orthogonal methods of detection: our previously published long-read sequencing SV callset derived from *de novo* assemblies for each strain compared to the GRCm39 reference genome [25], copy number (CN) estimates generated using QuicK-mer2 from Illumina short-read sequencing data [50], and SV calls determined with Bionano optical mapping [51]. High-confidence SVs were supported by two or more of our methods (Methods).

With this ensemble approach, we identify 146 deletions, 66 duplications, and 11 inversions for a total of 223 nonredundant, high confidence SD-mediated rearrangements across eight mouse strains. We find 40.5% of the (86/212) deletions and duplications were identified with QuicK-mer2, optical mapping, and at least one of the two long-read SV calling tools we utilized (Fig 4A).

**Figure 4.**
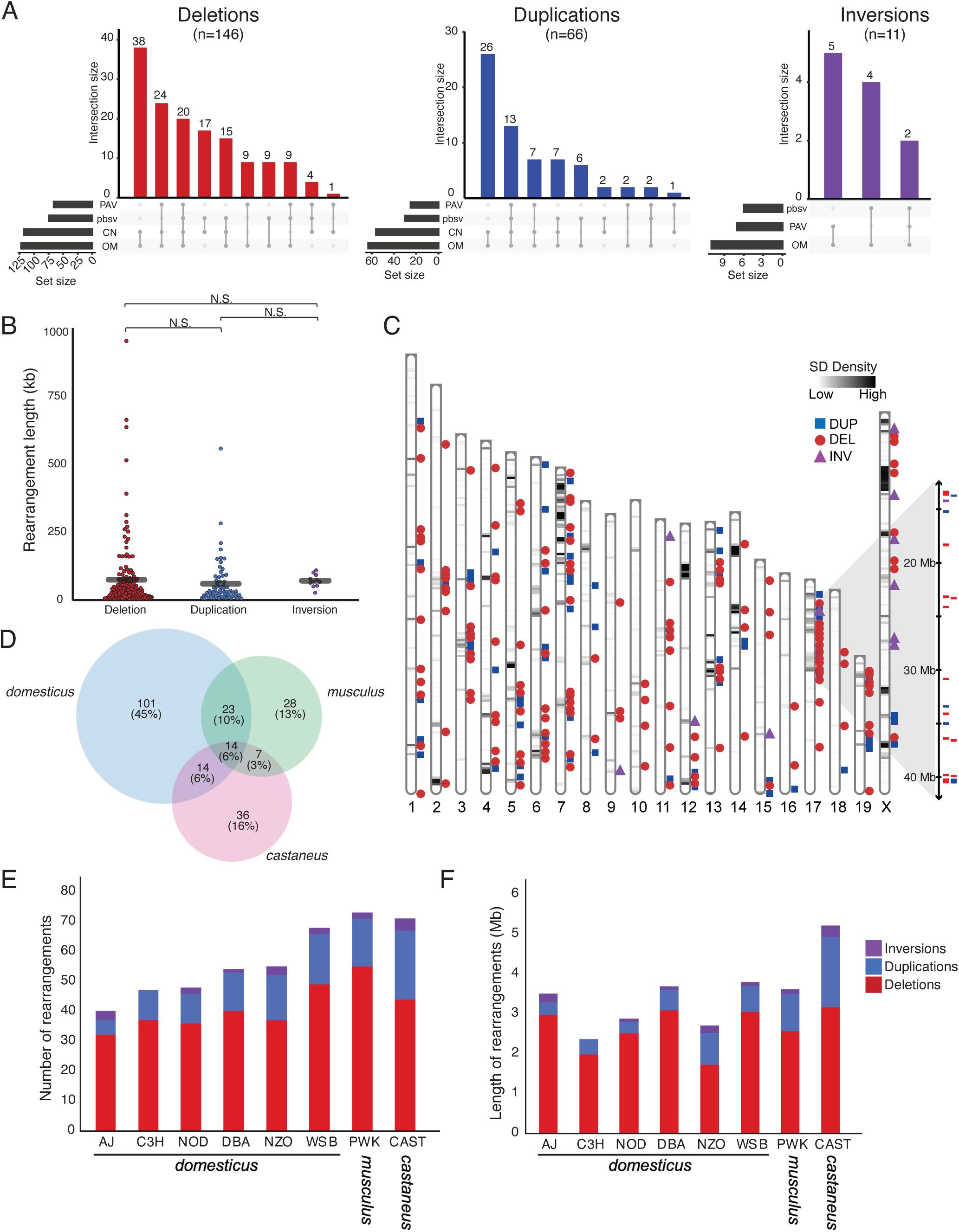
Identification of SD-mediated rearrangements in 8 strains of laboratory mice. (A) Total number of SD-mediated deletions (red), duplications (blue), and inversions (purple) identified by which orthogonal method: PAV, pbsv, optical mapping (OM), and copy number estimates from QuicK-mer2 (CN). (B) Length of SD-mediated rearrangements identified by variant type. (Mann Whitney U test) (C) SD-mediated rearrangement landscape in comparison to the SD density across the GRCm39 reference genome. Deletions (red circle) and duplications (blue square) are spread across most chromosomes. Inversions (purple triangle) occur most often on chromo-some X within the 8 strains examined. (D) Total number of SD-mediated rearrangements identified and shared between each subspecies of *Mus Musculus* (*domesticus*: A/J, C3H/HeJ, DBA/2J, NOD/ShiLtJ, NZO/HILtJ, WSB/EiJ; *musculus*: PWK/PhJ; *castaneus*: CAST/EiJ). (E) Count of SD-mediated rearrangements per strain. Rearrangements identified in multiple strains are counted for each strain. (F) Total length of all SD-mediated rearrangements per strain. Rearrangements identified in multiple strains are counted for each strain.

Although human SD-mediated rearrangements range from a few kb [7, 52, 53] to greater than 1 Mb [54–56], the 223 rearrangements we identify in mouse genomes are only up to ∼950 kb in length (Fig 4B), and many were less than 200 kb. The difference in size distributions may reflect challenges in assembling SD-rich regions or detecting large rearrangements between highly similar sequences [3, 57]. Alternatively, such large rearrangements may be less common in mice due to differences in SD organization [4], or because larger rearrangements are often disease-causing or deleterious [58], which have a greater impact in inbred mice where high homozygosity increases vulnerability to these mutations.

SD and rearrangement length are correlated (Supp 4A) and SD-mediated deletions and duplications are distributed across mouse genomes. However, a notable departure from this trend occurs on Chromosome 17 where within a 27 Mb hotspot (from ∼13 to ∼40 Mb) harbors 12 deletions, 5 duplications, and 1 inversion. This locus corresponds to the *t*-haplotype, a well characterized meiotic driver associated with transmission ratio distortion and male sterility in heterozygotes. The *t*-haplotype is made up of a series of overlapping inversions and other SVs that suppress recombination, but its architecture has remained poorly resolved due to short-read sequencing limitations. Previous studies have shown that the *t*-haplotype carries extensive copy number variation (CNV) and altered gene expression [59]. This Chromosome 17 region exhibits the highest density of SD-mediated rearrangements, despite having only a moderate density of SDs. In addition to this Chromosome 17 locus, more than half (6/11) of the identified inversions are present on the X Chromosome, which could be attributed to the high density of SD paralogs in inverted orientation (Fig 4C), a phenomenon also observed in humans [60].

Many SD-mediated rearrangements are subspecies-specific (165, 74%), while a smaller subset are shared between two or more subspecies (Fig 4D). These 58 shared variants are potentially indicative of recurrent SVs, errors in the GRCm39 reference, or variants in C57BL/6J [61]. 20 are present only in the wild-derived strains in our study (WSB/EiJ, PWK/PhJ, and CAST/EiJ) with three shared by all of these strains. The remaining 38 are present in many of the eight strains (Supp 4B), potentially reflecting recurrent or common rearrangements. We next sought to identify these 38 variants in C57BL/6NJ, a strain closely related to the reference strain, to estimate how many were indicative of reference errors or rearrangements in the B6J/NJ lineage. We detected 3 variants using PAV and/or pbsv in C57BL/6NJ suggesting that the remaining 35 variants likely reflect recurrence, false positives, or unresolved reference artifacts. However, interpretation is complicated by the mixed ancestry of laboratory mouse strains, many of which were derived from multiple subspecies during domestication and selected for desirable traits.

In total, these 223 rearrangements contribute to over 14.7 Mb of sequence variation between the 8 strains and GRCm39. Eight of the events occur as a deletion in some strains and a duplication in others, indicating recurrent reciprocal rearrangements at the same locus (Supp 4C-D). Strains more diverged from the reference genome, PWK/PhJ and CAST/EiJ, contain more variants and more base pairs altered by SD-mediated rearrangements than the strains in the *Mus musculus domesticus* lineage (Fig 4E-F).

To determine the accuracy of our SD-mediated rearrangement predictions, we next examined whether mouse SVs occur between SD paralogs not predicted by our criteria (Fig 3). Of the 223 rearrangements we identified 152 occurred between predicted SD paralogs. Two of the 152 rearrangements were deletions that intersect with genes predicted to result in lethality (*Pnp* and *Cdca8*), however, both only affect intronic or UTR regions of these genes. Notably, the remaining 71 rearrangements (32%) were mediated by SDs that failed our prediction criteria by having less than 95% identity in the reference genome (Supp 4E). To examine whether our predictions were valid for genomes with ∼0.5 million years of divergence from the reference, we investigated the identity of these 71 SD paralogs across our 8 strain assemblies.

For each of the 71 rearrangements mediated by SD paralogs with less than 95% identity in the reference genome, we examined the SD annotations generated for each strain (Methods). For SDs with less than 95% identity (<0.95 fracMatch) in GRCm39, the mean identity of SD paralogs in our strain assemblies show an increase by ∼0.12% (median: -0.0095%). For all SDs that mediate deletions including those with over 95% sequence identity we find that on average SDs within the samples are ∼0.17% more diverged (median: -0.00005%) than what is identified in the reference genome (Supp 4F). These findings suggest that while some rearrangements may involve SDs with lower identity than 95%, strain-specific mutations may alter paralog identity leading to SD pairs more (or less) capable of mediating rearrangements.

In light of this discrepancy, we wanted to further refine our predictions for loci likely to be affected by SD-mediated rearrangements between mouse genomes. Therefore, we developed a random forest machine learning model to identify SD pairs. The model was trained on a random selection of the SDs responsible for mediating mouse rearrangements and SD paralogs that were not involved in rearrangements within our cohort (Methods). The intervening length, or the distance between paralogs was the most influential parameter in determining the likelihood of SD-mediated rearrangements followed by the percent identity of SD paralogs. The length of the SDs and the orientation were less important factors contributing to SD rearrangements (Supp 4G). The model was then tested on the remaining SD paralogs involved in generating rearrangements and other randomly selected SD paralogs. Using a threshold of 0.6, the model achieved an F1 score of 84.6% (Supp 4H). We then determined that only 1,948 (13.2%) of the 14,773 predicted SD paralog pairs determined using parameters from human rearrangements were identified by our machine learning model. SDs predicted to undergo rearrangement based on the random forest model tended to have shorter intervening sequence lengths (∼23,140 bp) than those not predicted with our model (∼1,702,380 bp) (*P*-value <2.2e-16, Mann-Whitney U test) and have higher sequence identities (97.3% vs 96.7%) (*P*-value <2.2e-16, Mann-Whitney U test).

Overall, we find that large regions of the mouse genome are susceptible to SD-mediated variation, that this susceptibility likely contributes to ongoing genome instability and structural diversity across strains, and that intra-species SD-mediated variation in mouse genomes tends to comprise shorter rearrangements than those seen in between human genomes.

### Mouse diversity in gene copy number and expression

To identify how SD-mediated rearrangements might affect mouse phenotypes, we assessed protein classes subject to CNV and potential functional impacts of these changes. First, we used the PANTHER database to determine protein class families with CNV due to SD-mediated rearrangements (Methods) [62]. Overall, 215 genes were affected by SD-mediated copy number changes in the 8 mouse genomes; 148 genes intersect with deletions and 78 genes intersect with duplications. These genes include transcription factors, immune related genes, metabolite interconversion enzymes, and most abundantly transmembrane signal receptors (Fig 5A). For genes that intersect the breakpoints of inversions, proteins involved in storage and translation were often affected (Supp 5A).

**Figure 5.**
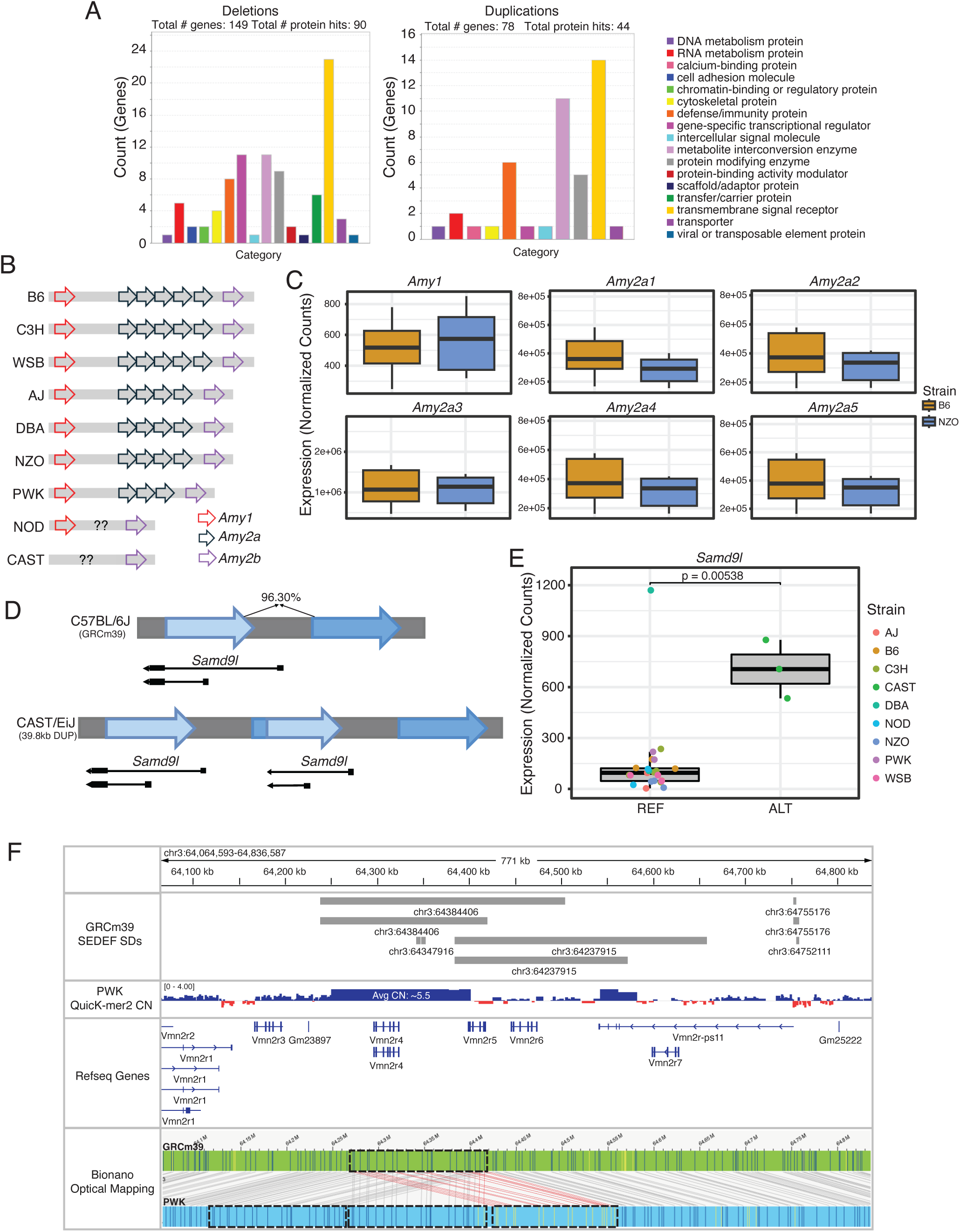
Genes affected by identified SD-mediated rearrangements. (A) PANTHER protein classes of genes that intersect with deletions (left) and duplications (right). (B) Representation of the copy number variation of Amy2a within Mus Musculus. (C) Amy2a gene copy expression differences between C57BL/6J (B6) and NZO/HILtJ (NZO). (D) Schematic of duplication of Samd9l in CAST/EiJ. (E) Samd9l expression in strains with reference allele (REF) and alternative allele (ALT). (F) Multi-copy duplication in PWK/PhJ as seen by copy number variation (QM2 CN) and optical mapping. Red lines were manually included for fluorescent labels that match in GRCm39 to PWK.

One locus of interest in adaptation to domestication is the amylase locus (AMY). Amylase enzymes digest starch into smaller molecules. Mammalian genomes often have both *AMY1*, the salivary amylase, and *AMY2a* and *AMY2b*, the pancreatic amylases. The AMY locus has variable copy number across mammals, often resulting in copy number amplification of these genes in domesticated species [13]. In mice, each copy of *Amy2a* in the GRCm39 reference genome is contained within an SD copy (Supp 5B). Previous studies in mice have identified *Amy2a* CNV between subspecies of *Mus musculus* [63]. Our data indicate that this variability also occurs within the *Mus musculus domesticus* subspecies, mirroring human variability at the *AMY1* locus [14, 15]. We manually reconstructed the mouse AMY locus using optical mapping data (Methods) for all 8 strains and found that C57BL/6J, C3H/HeJ and WSB/EiJ have 5 *Amy2a* copies, while the A/J, DBA/2J, and NZO/HILtJ strains have four. Additionally, we also confirmed the presence of three *Amy2a* copies in the PWK/PhJ strain (Fig 5B, Supp 5C) [63]. Notably, while *Amy1* and *Amy2b* were identified in NOD/ShiLtJ and *Amy1* was identified in CAST/EiJ, the *Amy2a* region could not be resolved in these strains with either optical mapping or our assemblies. While this does not mean these genes are absent from NOD/ShiLtJ and CAST/EiJ, we cannot definitively determine their presence or copy number. Mouse variation departs from the locus in the rat (*Rattus norvegicus*) genome, which has 1 copy of each of the amylase genes (*Amy1*, *Amy2a*, and *Amy2b*), and multiple proximal amylase pseudogenes [64]. To determine if *Amy2a* copy number difference resulted in expression changes, we next used RNA-seq data from pancreatic islet cells derived from C57BL/6J and NZO/HILtJ (Methods) (GEO:GSE235481). While we expect expression of *Amy2a1* to be significantly decreased in NZO/HILtJ compared to C57BL/6J, short read RNA sequencing data may not properly align reads to the correct copy of each *Amy2a* gene. As a result, we see a decrease in all copies of *Amy2a* in NZO/HILtJ compared to C57BL/6J (Fig 5C).

A 39.8 kb duplication of the *Samd9l* gene is present only in CAST/EiJ (Fig 5D). Samd9l is involved in proliferation and differentiation of cells, particularly hematopoietic stem cells, and is expressed in embryonic stem cells [65]. Regardless of where the breakpoint of the duplication resides within the SD, the duplicated copy of *Samd9l* is truncated in CAST/EiJ. We wanted to further investigate if there was an increase in expression in embryonic stem cells (Methods). We see that in CAST/EiJ, *Samd9l* is expressed approximately 7 times higher than in strains with the reference allele (*P*-value = 0.00538, Wald test) (Fig 5E). The increased expression may be due to the partial gene copy, or differential regulation of the locus [66–68]. We therefore examined the duplication of regulatory elements or changes in chromatin accessibility using Ensembl Variant Effect Predictor (VEP) (Methods) [69]. Our analysis indicates that the *Samd9l* duplication includes regulatory elements that could enhance gene expression, contributing to the observed increase in *Samd9l* expression. Additionally, the duplication encompasses the 5′ UTR and the transcription start site (TSS), which may further influence the elevated expression levels observed in CAST/EiJ. Deletion mutants and defective alleles of *Samd9l* in both humans and mice are known to disrupt hematopoiesis and can lead to bone marrow failure and leukemias [70, 71].

We also identified a multi-copy number variable rearrangement on Chromosome 3 that intersects with two vomeronasal 2 receptor genes, *Vmn2r4* and *Vmn2r5* in PWK/PhJ. QuicK-mer2 identifies an average copy number of 5.6 within the region where a triplication was identified from optical mapping data (Fig 5F). Vomeronasal receptor genes, amylase genes, and the *Samd9l* locus are all representative variants from our cohort of 126 SD-mediated variants that affect genes due to deletion, duplication, and inversion of loci that may impact fitness or disease predisposition.

## DISCUSSION

We provide a comprehensive analysis of SDs in the GRCm39 reference genome, data supporting how these duplicons arose, and define the role of SDs in rearrangements across diverse laboratory mice, contributing to our understanding of how SDs shape genomic variability within species. This work addresses a gap in mouse genomics, where SD regions are often excluded from analyses due to their repetitive nature and the difficulty of resolving them with short-read sequencing technologies [26, 72, 73]. However, SDs are far from genomic noise. They are drivers of genome evolution, structural variation, and functional diversity. Our SD annotation of nine mouse genomes elucidates both genome architecture and phenotypic variation across widely used laboratory mouse strains. These data can be further used to guide experimental design, allow for exclusion of these complex loci from certain analyses, and improve our understanding of gene content and the effect of variation in SD copy number and rearrangements. Our annotation of mouse SDs have also allowed us to investigate the organization of SDs within the mouse genome. While both our work and previous studies found that mouse SDs are more often tandem than human SDs [4], we find that mouse inverted paralogs are more dispersed than those in direct orientation, with a majority of inverted paralogs having >1 Mb of sequence between copies.

We identified the enrichment of both LINE-1 and ERV TEs at the flanks of mouse SDs, with LINE-1 elements being the most common. Our results indicate that both the age and the number of LINE-1 elements within a subfamily contribute to their ability to mediate homology directed repair leading to mouse SD formation. We observe no significant enrichment of the youngest LINE-1 subfamily, L1MdTf, which are active within the last 0-1 million years and are largely restricted to *Mus musculus domesticus* [25]. The lack of L1MdTf enrichment may therefore reflect its very recent expansion and limited evolutionary time to mediate rearrangements. The presence of LINE-1 and ERV elements at SD flanks indicates that they may be more likely to drive TE-mediated rearrangements in the mouse genome, contrasting with human TE-mediated rearrangements which are primarily driven by SINE *Alu* elements [41]. The mechanisms driving these rearrangements may also differ, as *Alu* sequences often lack substantial homology for NAHR, whereas LINE-1s are more likely to be substrates for homologous recombination. The TE differences in SD formation could underscore both the tandem structure of mouse SDs likely caused by homology/NAHR and the relatively more complex structure of primate and human SDs, which may have arisen through microhomology-mediated mechanisms associated with complex structural variant formation [10, 74].

We observed that TEs flanking SDs tend to exhibit greater sequence divergence than internal TEs. This may be caused by several factors. One possibility is that these flanking elements are chimeric, generated by past recombination events, such as NAHR, between divergent LINE-1 copies (Fig 2C). Additionally, flanking TEs have predated the duplication events, allowing more time for sequence divergence through accumulated mutations. In contrast, internal TEs were likely duplicated along with the SD and thus retain higher sequence identity. These findings suggest that TE composition and evolutionary history contribute not only to the formation of SDs but also to their subsequent sequence evolution and structural variation across strains.

We additionally developed a catalog of hotspots for potential SD-mediated rearrangements. With predictions, we can better anticipate SD-mediated rearrangements that may arise during the maintenance of mouse strains and mES cell lines, or anticipate *de novo* events in studies utilizing mouse models. In human genomes some loci have rates of rearrangement as high as 6.96x10^-5^ [75]. While some loci may not tolerate rearrangements due to the presence of essential genes, rearrangements in these loci may only intersect intronic regions or UTRs and can therefore be segregating in the population. We also determined that characteristics for predicting rearrangements in humans may need to be refined for other mammalian species or for non-pathogenic rearrangements.

To overcome the limitations of short-read sequencing in resolving complex, repetitive regions, we employed a multi-platform approach that leveraged the strengths of long-read sequencing, optical mapping, and short-read copy number estimation. In particular, long-read sequencing spanned large, repetitive elements with high resolution; optical mapping provided long-range, genome-wide structural context; and short-read data contributed reliable copy number estimates. With an ensemble approach we have identified 223 SD-mediated rearrangements across 8 diverse strains of inbred mice that contribute to over 14 Mb of genomic variation. While the rearrangements in mouse genomes are generally shorter than pathogenic rearrangements in humans [54, 55], this may be due to the organization [56] of SD in mouse genomes, the tendency for paralogs to be more tandemly duplicated, or the absence of disease-causing variants present in mouse genomes, as mice are bred in a highly controlled manner [4, 58]. While non-pathogenic CNVs in human genomes can be up to multiple megabases in length, they are rare events. On average non-pathogenic CNVs within the human population are ∼100 kb [76], which is comparable to the lengths of SD-mediated rearrangements in mouse genomes.

Amongst the 223 rearrangements, we identify 8 SD paralogs that mediated recurrent events, with some strains exhibiting deletions and others duplications at the same locus. These events highlight hotspots of SD-mediated rearrangements, consistent with observations from disease-associated regions such as the 15q11-q13 in human genomes, where recurrent deletions and duplications arise due to misalignment of SDs [16]. While many recurrent rearrangements have been identified at disease-associated loci, many neutral rearrangements (notably inversions) are also recurrent [77]. While these 8 loci represent clear examples of recurrent rearrangements, additional hotspots may exist but remain difficult to detect due to the challenges of accurately resolving breakpoints or long haplotypes within highly identical SD sequences.

The identification of rearrangements affecting gene copy numbers, such as the *Amy2a* locus and *Samd9l*, underscores the functional consequences of SD-mediated rearrangements. The variability in *Amy2a* copy numbers between strains demonstrates how SDs can drive copy number variation within a species, potentially leading to differences in traits such as starch metabolism. Similarly, the duplication of *Samd9l* in the CAST/EiJ strain, highlights how SDs can influence gene regulation and expression, potentially leading to strain-specific phenotypic differences. Previous research has also identified immune-related variation linked to SDs in mice [4, 23]; this is further supported by our findings, which reveal SD-mediated variation in copy number of immune-related genes (Fig 5A). While these strains have variable immune responses [78, 79], these differences cannot be directly linked to the variation we identified and would require more targeted studies for further investigation. In addition, our results demonstrate how SDs contribute to the diversity of the amylase gene family and olfactory receptors, illustrating the potential impact of SDs on metabolic processes and sensory adaptation. Future work incorporating long-read RNA sequencing approaches could provide deeper insights into the RNA expression profiles and isoform structures of these variants. Additionally, long-read RNA characterization could help resolve expression differences between highly similar gene copies, such as those within the amylase locus, by capturing full-length isoforms and identifying sequence differences that distinguish paralogs.

Our study has enabled future investigation of mechanisms of SD-mediated rearrangements within the mouse genome by providing a robust catalog of reference SDs and a thorough characterization of variants across inbred mouse strains. Additionally, our data provides a foundation for comparing mechanisms of SD-mediated rearrangements in mice to those observed in human and other mammalian genomes, with particular interest in the role of PRDM9 binding sites, non-allelic homologous recombination (NAHR), and other potential mechanisms driving genomic instability. While we observe that SD paralogs sometimes have lower overall sequence identity than expected, localized regions of high homology may still exist between paralogs, potentially sufficient enough to mediate rearrangements through NAHR. This remains an important area for future study. Our data offer a valuable resource for cross-species comparisons of SD formation and rearrangement. By elucidating the role of TEs, recombination, and genomic architecture in shaping structural diversity, our findings highlight the importance of SDs as dynamic elements that drive genome evolution, phenotypic variation, and potentially, adaptation. This work represents an essential step toward understanding the interplay between repetitive elements, genome structure, and evolution in a key biomedical model system.

## LIMITATIONS

The remaining limitations of this study include the resolution of highly repetitive loci, particularly on Chromosomes 7 and the X Chromosome. Improved methods of genome sequencing and assembly now exist that use a combination of PacBio HiFi and Oxford Nanopore Technologies ultra long reads to allow for the generation of highly accurate, near-T2T genome assemblies that can better capture these long SDs and other highly repetitive loci [3]. In addition, we were unable to identify rearrangements on the Y Chromosome for multiple reasons. First, due to technical limitations of SEDEF, the highly repetitive nature of the Y Chromosome resulted in failed annotation. Additionally, the long-read sequencing and genome assemblies for these 8 strains was performed on female mice and therefore we did not assemble the Y Chromosome. To address this limitation, more contiguous assemblies for male mice would be important. This will allow us to capture the repeat-rich Y Chromosome and significantly improve our ability to capture and accurately map SD-rich regions.

Additionally, our genome assemblies were generated without phasing. While laboratory mice are highly inbred, ∼5% of their genomes remain heterozygous [80]. Although this limitation is reduced in our study, as we examined copy number estimates generated with QuicK-mer2 to infer potential heterozygous SD-mediated rearrangements, phased genomes would be important for examining the most rapidly evolving loci in this model organism. Moreover, with haplotype-resolved, complete assemblies, we would be less likely to have false negatives in our call set and could more accurately determine false positive SD-mediated rearrangement call rates.

Finally, while our study focused on standard laboratory mouse strains, future work will benefit from including more genetically diverse mice from different species to explore the broader spectrum of SD-mediated rearrangements. The SPRET/EiJ (*M. spretus)* strain is more distantly related (∼2 million years diverged) to the strains utilized in this study and therefore is likely to have rearrangements that are absent or less common in inbred mouse strains, offering valuable insights into the evolutionary dynamics of SDs and their role in species divergence. Examining these rearrangements in more genetically diverse models can help elucidate how SDs have contributed to adaptive evolution, species-specific traits, and genomic instability. This would also allow us to better understand the evolutionary history of SDs in shaping genomic architecture within a species.

## METHODS

### Annotation of segmental duplications in the mouse genome

The set of SDs was generated on the GRCm38 (mm10) reference genomes using SEDEF (v1.1-35) and BISER (v1.4) [33, 81]. SEDEF annotations were additionally generated on GRCm39. Soft masked reference genomes for GRCm38 and GRCm39 were obtained from the UCSC genome browser and the Y Chromosome was removed from each of the reference genomes prior to running SEDEF due to technical issues. Post annotation, SDs with over 50% tandem repeat composition using bedtools (v 2.31.1) to intersect SD and simple repeat data from tandem repeats finder (TRF) extracted from the UCSC genome browser [82]. The filtered output files from SEDEF, which originally contains one row per SD pair (with one SD designated as the primary and its paralog as the secondary), was expanded so that each SD had its own row. This transformation facilitates downstream analyses by allowing each SD to be treated as an independent entry while still retaining paralog information. SD annotations generated by SEDEF were then filtered for length ≥ 1 kb and mismatch rate (X) ≤ 10.

To allow for consistent comparisons in GRCm38 to establish the best SD annotation method, SDs identified by BISER present on the Y Chromosome and SDs with over 50% tandem repeat content were removed from the annotation set, similar to the method used for generating a SD annotation set using SEDEF. WGAC SDs for GRCm38 were extracted from UCSC table browser [5, 83] and were also filtered to remove SDs present on the Y Chromosome.

Comparisons between length and percent identity of SDs identified by WGAC, BISER, and SEDEF was completed in R (v4.3.1). We compared all SD annotations identified by each tool as well as only those annotated as intrachromosomal. We additionally examined the overlap of unique basepairs from each tool. SDs from each tool were first merged into unique regions using bedtools merge (v2.31.1). These unique regions of SD were then used to identify and quantify overlaps and unique regions between the three tools using GenomicRanges (v1.16.0) in R (v4.3.1). The same analysis of overlapping SD regions for intrachromosomal SDs was completed using the same method.

### Sample selection, sequencing, and variant calling

Long-read sequencing genome assemblies and structural variant (SV) callsets for eight laboratory mouse strains (A/J, CAST/EiJ, NOD/ShiLtJ, NZO/HILtJ, PWK/PhJ, WSB/EiJ, DBA/2J, and C3H/HeJ) were utilized from previously published data [25]. These strains were chosen for their high genetic diversity [25], and many are the founders of the collaborative cross. PAV duplication calls were generated by SV-Pop [26]. Briefly, SV insertion sequences were mapped back to the GRCm39 reference genome with minimap2 [84] with parameters “-x asm20 -H –secondary=no -r 2k -Y -a –eqx -L -t 4” and selecting aligned records indicating the duplicated reference loci for mappable insertion sequences.

### Copy number estimation by QuicK-mer2

Previously published Illumina short-read sequencing data [23] was used to examine copy number variation using QuicK-mer2 (v2021-04-28) [50]. QuicKmer2 search was first run on GRCm39 to generate a unique set of k-mers. The -c option was used to specify known copy number stable regions. We used the toInclude.bed file that was generated for GRCm38 (https://kiddlabshare.med.umich.edu/QuicK-mer/QuicK-mer2-refs/mm10/ref/) and lifted over to GRCm39 coordinates using Liftover (UCSC). We then ran quicKmer2 count to calculate the occurrences of each k-mer in our eight samples and quicKmer2 est to correct values based on GC content and generate copy number estimates in 1 kb windows across the genome.

### Bionano optical genome mapping

Large genomic rearrangements were identified using Bionano optical mapping [51, 85]. Frozen mESC lines from the eight strains were submitted to Bionano Genomics for processing (∼3x10^6^ cells per strain). Bionano Genomics performed DNA extraction, labeling, and analysis using the Saphyr system. *De novo* and rare variant detection was conducted against the GRCm39 reference genome by Bionano Genomics and was further analyzed using the Bionano Access web server. Molecule coverage ranged from 317 to 496 molecules per genome position, with average molecule lengths between 68 kb and 111 kb. We utilized the vcf files generated by Bionano Genomics to examine potential SD-mediated rearrangements. Additionally, the labeling patterns were manually reconstructed on a case-by-case basis to establish SV structure by comparing the sample assembly and molecule labeling pattern to the labeling pattern of the reference.

### Annotation of segmental duplications in long-read genome assemblies

Segmental duplications were also annotated in previously published long-read genome assemblies for the eight mouse strains. Assemblies were soft-masked using RepeatMasker (v4.1.2) (https://www.repeatmasker.org/) prior to SD annotation using SEDEF [33]. The raw SD annotations were processed using the same filtering criteria as described for the reference genome annotations. This discrepancy in the number of SDs annotated in these assemblies compared to GRCm39 likely reflects a combination of assembly collapses or difficulty resolving structurally complex repetitive loci. The genome assemblies have N50s averaging ∼8 Mb, and consist of ∼2,340 contigs per strain. Interestingly, while PWK/PhJ has the highest number of SDs, it also shows the least amount of SD base pairs among the strains. This indicates a larger number of shorter SDs within its assembly, which results in a higher concentration of SDs near the median size, in contrast to the more dispersed size distribution observed in other strains. The shorter length of SDs in PWK/PhJ may be due to a greater abundance of shorter contigs within the assembly and a lower N50, leading to fragmented assemblies, particularly within SD-rich regions as well as being more diverged from the reference genome. This could also be due to other factors not considered.

### Intersecting SD flanks with repetitive elements

To examine SD flanking regions for repetitive elements, we analyzed the SDs we annotated with SEDEF [33] in the mouse reference genome (GRCm39). Flanking regions were defined as 5 bp upstream and downstream of the SD start and stop positions. Overlapping flanking regions were merged into single continuous region usings bedtools (v2.31.1). These merged regions were then intersected with repetitive elements annotated in the mouse reference genome using the UCSC RepeatMasker track for GRCm39, which includes transposable elements and other repetitive elements.

We performed 1,000 permutations of the flanking regions using the regioneR package (v1.36.0) in R (v4.3.1), applying the circularRandomizeRegions function to preserve the spatial relationships of the flanking regions. The observed number of intersections between the original flanking regions and repetitive elements was compared to the mean of the intersections obtained across the 1,000 permutations. Statistical significance was determined by the regioneR package as an empirical P-value, defined as the proportion of permuted overlaps greater than or equal to the observed number of overlaps. We used the UCSC RepeatMasker track from GRCm39 to count the number of elements for the different LINE-1 subtypes as well as examine lengths of these events using R (v4.3.1).

### Estimating genetic distance between LINE-1 elements flanking and internal to SDs

To estimate the genetic distance of LINE-1s associated with SDs, we analyzed both flanking and internal LINE-1 sequences. For SD paralogs annotated as direct, LINE-1 sequences at the 5′ flanks were extracted from the GRCm39 genome using samtools (v1.21). These sequences were aligned pairwise using ClustalW (v2.1), and genetic distances were calculated using Kimura’s two-parameter model as implemented in the ape package (v5.8) in R (v4.3.1). A similar analysis was performed for LINE-1 elements located at the 3′ flanks of SD paralogs.

To investigate internal LINE-1s, we extracted sequences of LINE-1s present within SD paralogs that had matching LINE-1s at either the 5′ or 3′ flanks. Only internal LINE-1 elements meeting the following criteria were included: LINE-1 lengths were within 90% of each other, distances from the start position of the SDs were within 90% of each other, and LINE-1s belong to the same subtype (e.g., both L1MdA or both LX).

Statistical comparison of genetic divergence between flanking and internal LINE-1s was conducted using Wilcoxon rank-sum tests for unpaired comparisons and Wilcoxon signed-rank tests for paired comparisons within SD paralogs. Enrichment of specific LINE-1 subtypes at SD flanks were assessed using Fisher’s exact test. All tests were two-sided, and significance was considered at *P*-value < 0.05.

### Prediction of SD-mediated rearrangements

To predict SD-mediated rearrangements, we first excluded SD sequences identified by SEDEF that were located on the Y Chromosome, random chromosomal fragments, or unclassified chromosomal regions to ensure our analysis focused on well-characterized genomic regions. We then filtered for intrachromosomal SD paralogs with over 95% identity (X ≤ 5), longer than 1 kb, and having less than 10 Mb of intervening sequence between paralogs. These criteria were established based on known rearrangements in human and mouse genomes, as well as previous predictions of hotspots identified in human genomes.

Predicted SD-mediated rearrangements were characterized based on orientation of SD paralogs (direct and inverted) separately and examined characteristics of predicted SDs and loci. To assess whether the distance between predicted SD paralogs differed by orientation, we performed Welch’s two-sample t-test, which accommodates unequal variances between groups. This test compared the mean intervening sequencing lengths separating direct versus inverted SD paralogs.

Further characterization of predicted SD loci includes calculating the proportion of the genome flanked by these paralogs on each chromosome using GenomicRanges (v1.16.0) in R (v4.3.1). We calculated the unique basepair coverage predicted regions and divided by chromosome or whole genome length.

### SD-mediated rearrangement identification

To identify SD-mediated rearrangements, we utilized long-read SV calls from PAV and pbsv that were previously published [25]. We intersected these SV calls with SD annotations using bedtools (v2.31.1) intersect for direct intersection. We further filtered the SVs to include only those that intersected with an SD paralog pair at the 3′ and 5′ ends of the SV. For SVs that intersected with multiple SD paralogs, the pair with the highest identity was selected.

We used the vcf files containing SV calls generated from optical mapping in the same manner as SV calls from long-read sequencing approaches. In addition, we manually investigated the labeling pattern and molecules from the samples on Bionano Access. Labeling patterns indicating deletions and duplications between SD paralogs were used to determine SV lengths in these loci.

For short-read data, we used QuicKmer2 (v2021-04-28) to estimate copy number (CN) as described above [23, 50]. We examined the mean CN within the SV regions identified by PAV, pbsv, and Bionano optical mapping by taking the sum of the CN estimates for each base pair divided within the SV by the length of the SV. We classified deletions with a CN estimate of < 1.3 (1 for heterozygous or 0 for homozygous deletions) and duplications with a CN estimate of > 2.7 (3 for a heterozygous duplication, 4 for a homozygous duplication, or more for multi-copy regions). CN neutral regions have a CN estimate of ∼2 (CN ranging from 1.3-2.7).

Finally, we filtered for SD-mediated rearrangements using an ensemble approach, identifying those detected by at least two out of three methods: long-read sequencing SVs (grouped as one method), Bionano optical mapping, and QuicK-mer2. Rearrangements identified by different methods were merged if they occurred between the same SD paralogs, regardless of reciprocal overlap, due to variability in breakpoint resolution across platforms. For merging rearrangements across strains, we required that events involve the same SD paralogs and have a reciprocal overlap of >30%. Inversions relied on long-read and optical mapping SV calls, as the rearrangements would be copy number neutral.

We examined genome distribution of SDs and SD-mediated rearrangements using RIdeogram (v0.2.2). We compiled the SD density per Mb by calculating the total number of unique SD basepairs within each 1 Mb bin. SV types were clustered separately to examine differences in their distribution.

### Comparison of percent identity of SDs within the reference to samples

To compare the perfect identity of SDs between the reference genome and those in our individual assemblies, we first aligned the assemblies to the GRCm39 reference with minimap2 [25] to establish location of assembly SDs within the reference genome coordinates. We then sought to establish if the SV was contained within a single alignment record to the reference genome. This ensures that each SV occurred in a reliably mapped region in the assembly. We then matched the reference SDs to SDs in the individual assemblies by comparing their positions and requiring that the lengths of the sample and reference SDs to be within 10% of each other. Only SD pairs that had a single unique match between the reference and sample were retained. For each retained pair, we calculated the difference in percent identity (percent identity in the reference minus percent identity in the sample). SDs in the reference were categorized based on whether their percent identity was ≥95% identity or <95%, and these categories were compared to the percent identity observed in the corresponding SDs in the sample assemblies.

### Developing and using a model for predicting SD-mediated rearrangements

We utilized a random forest machine learning model using the randomForest package (v4.7-1.2) in R (v4.3.1) to predict whether SD paralogs will undergo rearrangement. The model was trained using the identified SD-mediated rearrangements and randomly selected SD paralogs within the reference genome using the length of the sequence between SD paralogs, SD length for both paralogs, percent identity between SD paralogs, and the orientation of SD paralogs. The model was trained on a random selection of 166 of the identified rearrangements and 249 random SD paralogs.

The model was then tested on the remaining identified rearrangements (56) and 84 random SD paralogs. To optimize the model, we tested multiple thresholds. A threshold determines the decision boundary for classification, influencing the balance between precision and recall. We selected a value of 0.6 based on the highest F1 score with the best tradeoff of precision and recall.

We then ran this model with a threshold of 0.6 on the 14,773 SD paralogs we predicted with our predetermined characteristics in order to identify SD paralogs likely to undergo rearrangements. We determined which predicted paralogs were more likely to undergo rearrangement based on intervening sequence length, SD lengths, percent identity (FRACMATCH), and orientation. Significant differences in intervening length and percent identity between those likely to undergo rearrangement based on the model and those not, was determined by the Mann-Whitney U test.

### Protein class analysis of genes affected by predicted and identified rearrangements

We examined the intervening regions of SDs that have the potential to recombine in the GRCm39 reference genome. These regions were intersected with the GRCm39 RefSeq using bedtools (v2.31.1) intersect and similar methods as the genes in predicted intervening regions [86]. We examined all genes within these regions, including only protein-coding genes (NM genes).

We then ran PANTHER pathway protein class analysis using the web-based interface of the PANTHER Classification System [62] on the genes within the intervening regions of predicted SDs in direct orientation or intersecting with the predicted inverted SD sequences. Lethality was assessed using MGI batch data analysis by examining the mammalian phenotype [87] (https://www.informatics.jax.org); any gene associated with a lethal phenotype in pre- or postnatal timepoints was binned as potentially lethal. The same analysis was performed on the deletions and duplications identified in the 8 strains in our study to examine which protein classes were most often affected.

We additionally examined the SD-mediated rearrangements that occurred in our strains for those that affected genes demarcated as potentially causing lethality. We identified 2 deletions that intersect with these genes and further examined their effects using the Ensembl Variant Effect Predictor (VEP) [69]. We additionally identified 6 duplications intersecting with these genes that have the potential to cause lethality, however they would not render the gene nonfunctional, which therefore may explain their presence within the strains. Other than examining lethality, we used VEP to examine the effects of some gene duplications, including *Samd9l* to establish how our variants may affect regulatory regions or coding sequences.

### RNA-seq analysis of expression of genes affected by SD-mediated rearrangements

We leveraged previously published RNA sequencing data from mESCs for these strains [25]. We utilized the available count tables for this dataset. Differential expression analysis of genes affected by SD-mediated rearrangements was performed using the same methods described in the original study using DESeq2 (v1.49.1) in R, which applies the Wald test for statistical significance and controls for multiple testing with the Benjamini-Hochberg procedure to adjust P-values.

We additionally leveraged publicly available pancreatic islet RNA sequencing count matrix tables for NZO/HILtJ and C57BL/6J (GEO:GSE235481) [88] to examine the Amylase gene cluster in these strains using DESeq2 (v1.49.1). We utilized only the control NZO/HILtJ (n=7) and C57BL/6J (n=8) mice (10% fat diet) for our analysis. P-values and adjusted P-values of expression differences between NZO/HILtJ and C57BL/6J for each gene analyzed were determined using DESeq2.

## Quantification and statistical analysis

Statistical tests were performed with R (v4.3.1).

## Author Contributions

C.R.B. conceived of, supervised, and procured funding for the study. E.R.F. performed all analysis, interpreted the results, and handled the data deposition. P.A.A. contributed to analytical discussions and data interpretation. A.F. gave critical feedback on analysis. P.B. assisted with data interpretation. E.R.F. and C.R.B. wrote the manuscript and P.A.A., A.F., and P.B. reviewed and edited the manuscript.

## Acknowledgements

This research was supported by The Jackson Laboratory Director’s Innovation Fund, The Jackson Laboratory Cancer Center (P30CA034196), and the National Institutes of Health (R35GM133600 to C.R.B. and T32HG010463 for support of E.R.F.). We would like to thank Anne Czechanski, Whitney Martin, and Laura Reinholdt for providing frozen cell stocks and facilitating submission to Bionano Genomics. We would also like to thank Beth Dumont for her review and critical proofreading of the manuscript. RNA-seq results for the amylase locus were obtained from NCBI (Bioproject PRJNA986136) and were generated with JAX institutional funds and work from Candice Baker, Isabela Gerdes Gyuicza, and Gary Churchill. Supplemental Figure 3A was created with the help of BioRender.

## Data and Code Availability

The segmental duplication annotations for GRCm39 and long-read *de novo* assemblies, QuicK-mer2 copy number estimates in bed and browser formats, and Bionano optical mapping assemblies for eight strains of inbred mice can be obtained through Zenodo at DOI: 10.5281/zenodo.15723435. All software used in this study can be found in the methods.

