## Supplementary material for "Segmental duplication-mediated rearrangements alter the landscape of mouse genomes": Supp 1

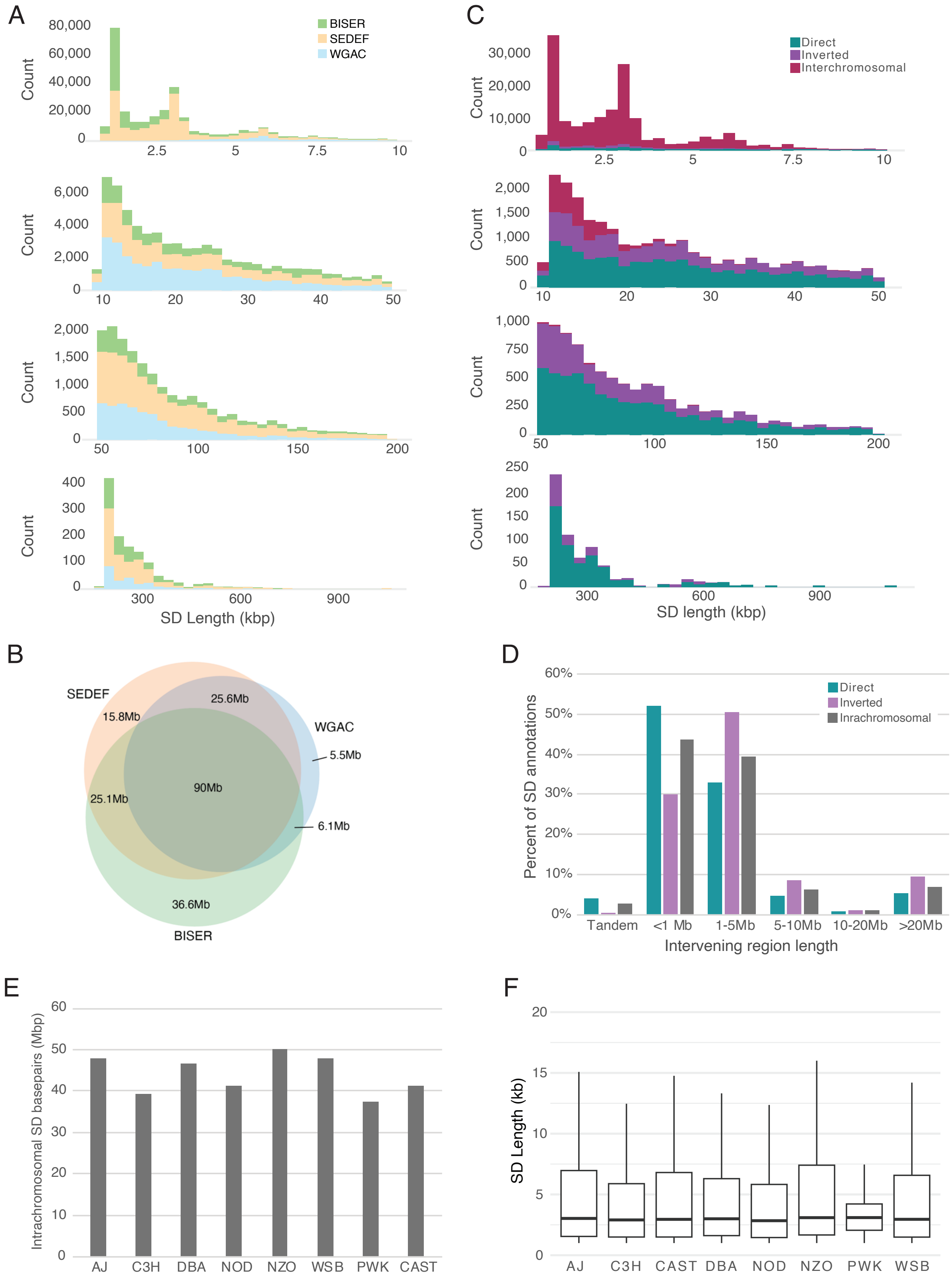
A
BISER
SEDEF
WGAC
Count
80,000
60,000
40,000
20,000
0
2.5
5
7.5
10
6,000
4,000
2,000
10
20
30
40
50
2,000
1,500
1,000
500
100
150
200
400
300
200
100
300
600
900
SD Length (kbp)
C
Direct
Inverted
Interchromosomal
30,000
20,000
10,000
750
250
SD length (kbp)
B
SEDEF
WGAC
BISER
15.8Mb
25.6Mb
5.5Mb
90Mb
25.1Mb
6.1Mb
36.6Mb
D
Direct
Inverted
Inrachromosomal
Percent of SD annotations
60%
50%
40%
30%
20%
10%
0%
Tandem
<1 Mb
1-5Mb
5-10Mb
10-20Mb
>20Mb
Intervening region length
E
Intrachromosomal SD basepairs (Mbp)
60
50
40
30
20
10
0
AJ
C3H
DBA
NOD
NZO
WSB
PWK
CAST
F
SD Length (kb)
20
15
10
5
0
AJ
C3H
CAST
DBA
NOD
NZO
PWK
WSB

**Supplementary figure 1** (A) Length distribution of SD annotations from BISER (green), SEDEF (yellow), and WGAC (blue) in GRCm38. (B) Base pair overlap of SDs identified by BISER, SEDEF, and WGAC in GRCm38. The Venn diagram shows the amount of SD sequence (in megabases) shared or uniquely identified by each method. The central overlap (90 Mb) represents regions detected by all three tools. (C) Length distribution of SD annotations in direct (teal), inverted (purple), and interchromosomal (red) in GRCm39. A majority of interchromosomal SDs were found to be shorter in length, with 97.5% of interchromosomal SDs being less than 10 kb while 40.7% and 46.2% of direct and inverted SDs are less than 10 kb in length. (D) Distribution of the distance between (intervening region) direct and inverted SD paralogs for SDs >10kb in length in GRCm39. (E) Non-redundant intrachromosomal SD basepairs annotated by SEDEF in eight mouse strains: A/J (AJ), C3H/HeJ (C3H), DBA/2J (DBA), NOD/ShiLtJ (NOD), NZO/HILtJ (NZO), WSB/EiJ (WSB), PWK/PhJ (PWK), and CAST/EiJ (CAST). (F) Lengths of SDs annotated in the eight mouse genomes in kilobases (kb).
