## Supplementary material for "Segmental duplication-mediated rearrangements alter the landscape of mouse genomes": Supp 2

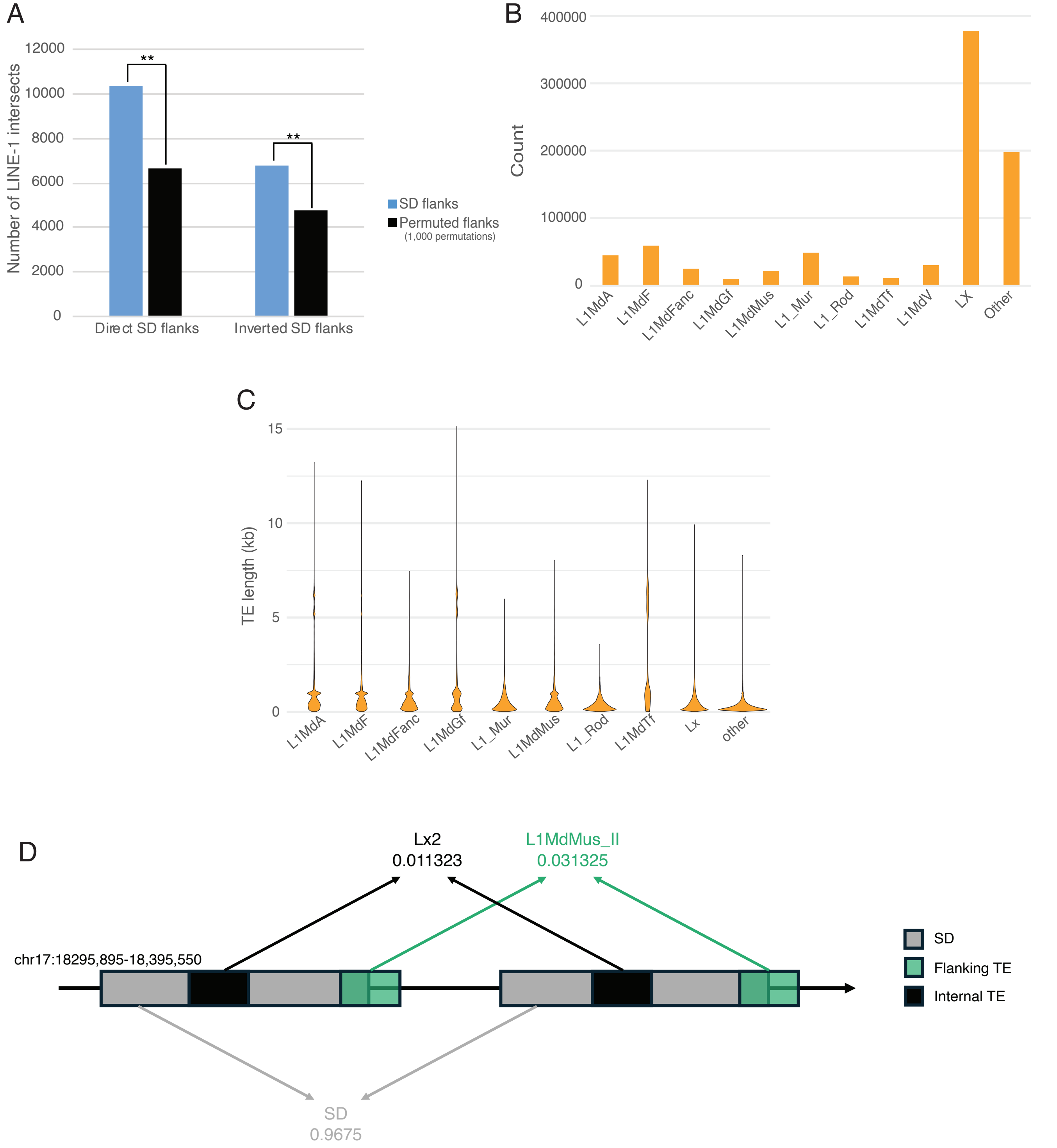


**Supplementary figure 2** (A) Number of intersects of direct and inverted SD flanks with LINE-1 elements compared with 1,000 permutations. (** = p ≤ 0.001). (B) Count of elements in different LINE-1 subtypes in GRCm39. (C) Length distribution of LINE-1 subtypes in GRCm39. (D) Schematic of genetic distance of TEs flanking and internal to SD paralogs.
