## Supplementary material for "Segmental duplication-mediated rearrangements alter the landscape of mouse genomes": Supp 3

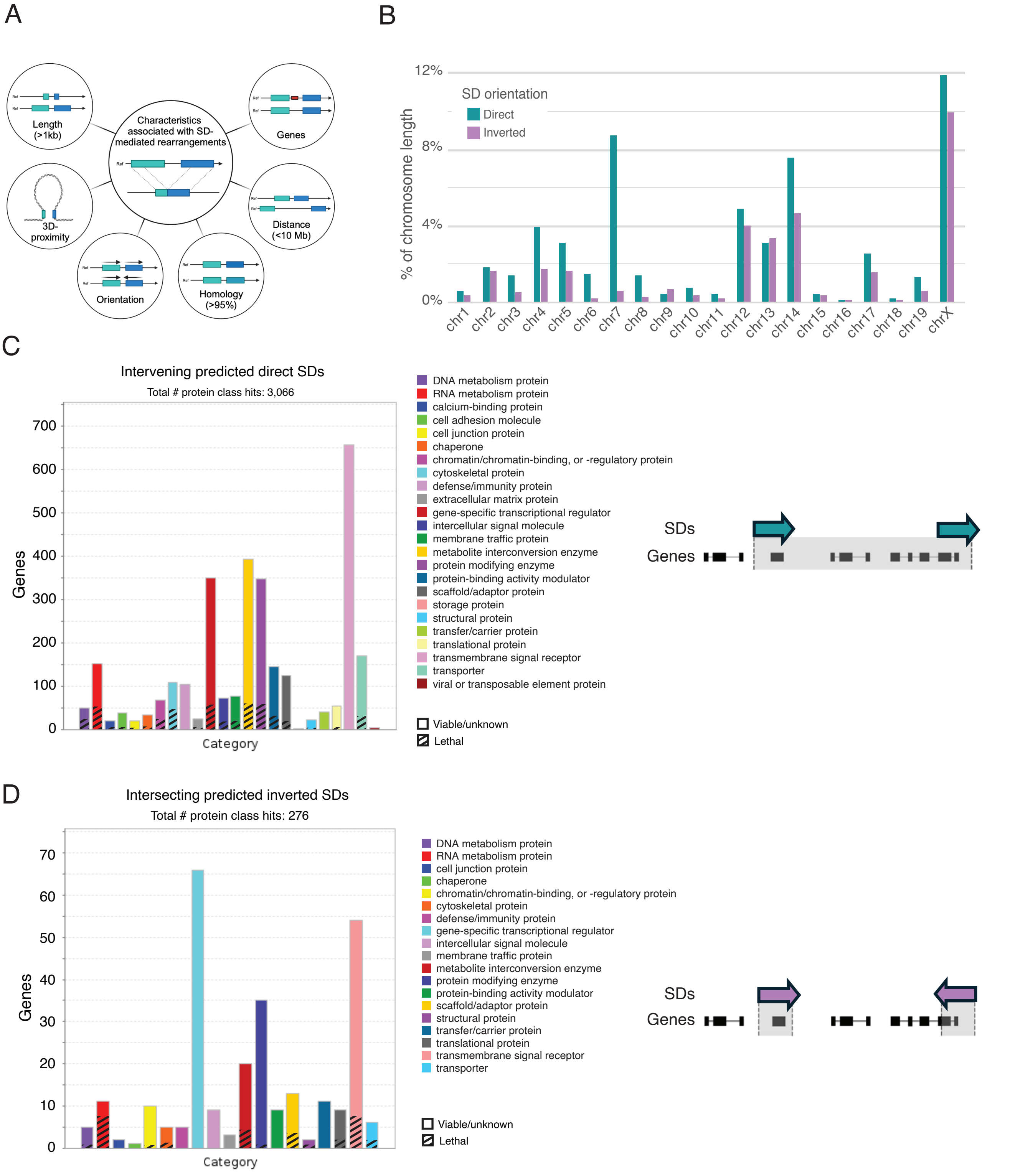


**Supplementary figure 3** (A) Characteristics that are associated with the likelihood that SDs will recombine. (B) Predicted SD composition of each chromosome (unique SD basepairs / chromosome length). (C) PANTHER protein classes of genes that reside within regions of the genome flanked by direct SD paralogs. (D) PANTHER protein classes of genes that reside within inverted SDs and have the potential to be disrupted by an inversion breakpoint.
