## Supplementary material for "Segmental duplication-mediated rearrangements alter the landscape of mouse genomes": Supp 4

A B

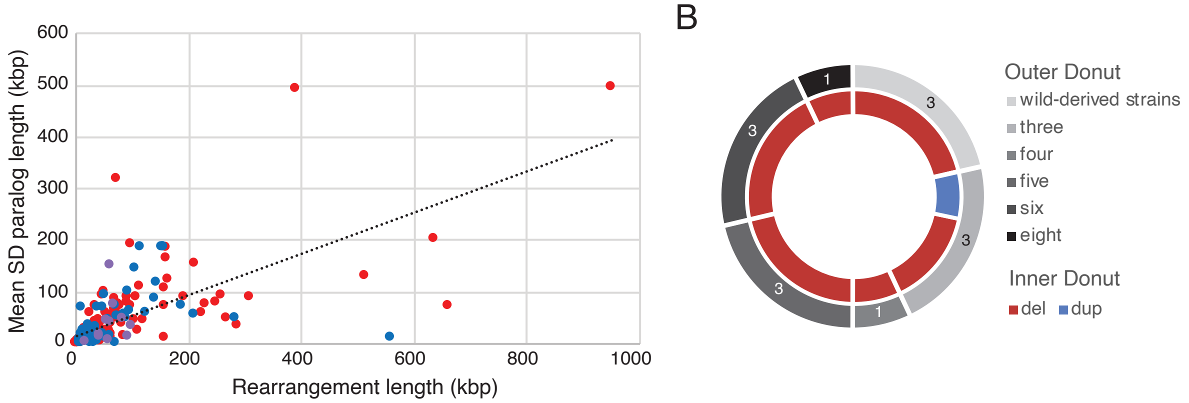


C

| Chr. | Start | Stop | Del. | Dup. | Size (kb) |
|---|---|---|---|---|---|
| chr1 | 31,303,658 | 31,311,212 | PWK | CAST | 7 |
| chr1 | 93,817,942 | 93,886,592 | AJ, C3H, DBA, NOD, NZO, PWK, WSB | CAST | 68 |
| chr3 | 88,274,796 | 88,289,684 | PWK | WSB | 14 |
| chr6 | 85,721,205 | 85,767,802 | PWK, WSB | NZO | 46 |
| chr7 | 106,704,514 | 106,727,043 | AJ, C3H, DBA, NZO | CAST | 22 |
| chr11 | 116,630,045 | 116,656,714 | AJ, C3H, DBA, PWK | NZO | 26 |
| chr13 | 22,886,856 | 22,953,240 | CAST | WSB | 66 |
| chr17 | 40,235,733 | 40,502,890 | WSB | PWK | 267 |

D

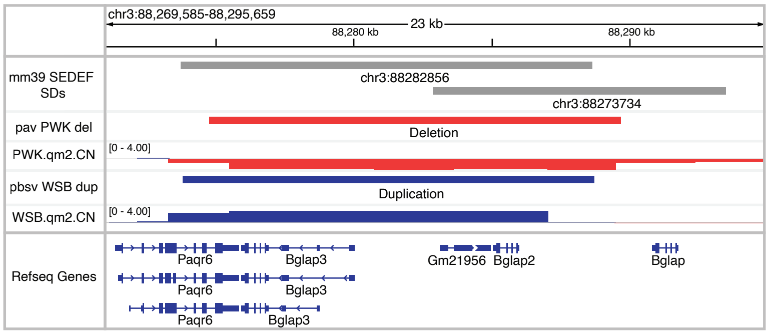


E F

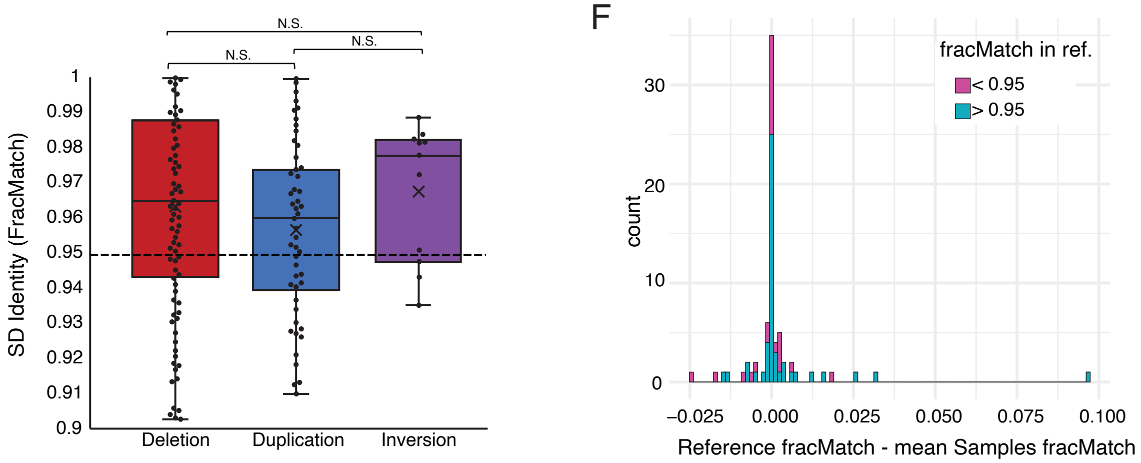


G H

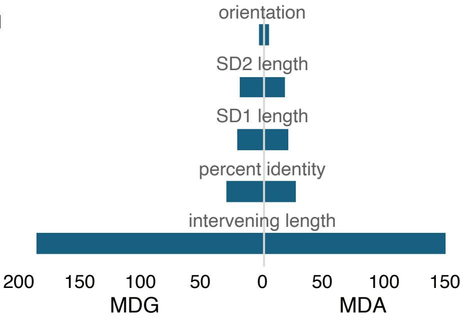


| Threshold | 0.7 | 0.6 | 0.5 | 0.4 | 0.3 |
|---|---|---|---|---|---|
| **Accuracy** | 0.9147 | 0.9147 | 0.8923 | 0.8878 | 0.8878 |
| **Precison** | 0.8135 | 0.7761 | 0.7222 | 0.7066 | 0.6962 |
| **Recall** | 0.8571 | 0.9285 | 0.9285 | 0.9464 | 0.9821 |
| **F1** | 0.8347 | **0.8455** | 0.8125 | 0.8091 | 0.8148 |

**Supplementary figure 4** (A) Rearrangement length compared to mean length of SD paralogs involved. (B) Number of strains where variants shared by all 3 subspecies were identified (outer donut) and whether these variants were deletions or duplications (inner donut). (C) Table of duplications and deletions mediated by the same SD paralogs in different strains. (D) Recurrent rearrangement of a deletion in PWK and a duplication in WSB mediated by the same SD paralogs. (E) Paralog identity of SDs that mediate deletions, duplications, and inversions. Points represent paralog identities for each rearrangement identified (Mann Whitney U test) (F) Difference in the fracMatch of reference SD paralogs that mediate deletions compared to the mean of the samples where the same SD paralogs were identified (values >0 indicate high SD percent identity in reference than in sample). SD paralogs with <95% identity in the reference shown in purple, SD paralogs with >95% identity in the reference shown in blue. (G) Characteristics included in the random forest model for predicting SD-mediated rearrangements. The Mean Decrease in Accuracy (MDA) and Mean Decrease in Gini (MDG) measure the importance of each feature in the model. Features  are ranked based on their contributions to model performance. (H) Metrics for assessing performance of the model using different threshold values.
