## Supplementary material for "Segmental duplication-mediated rearrangements alter the landscape of mouse genomes": Supp 5

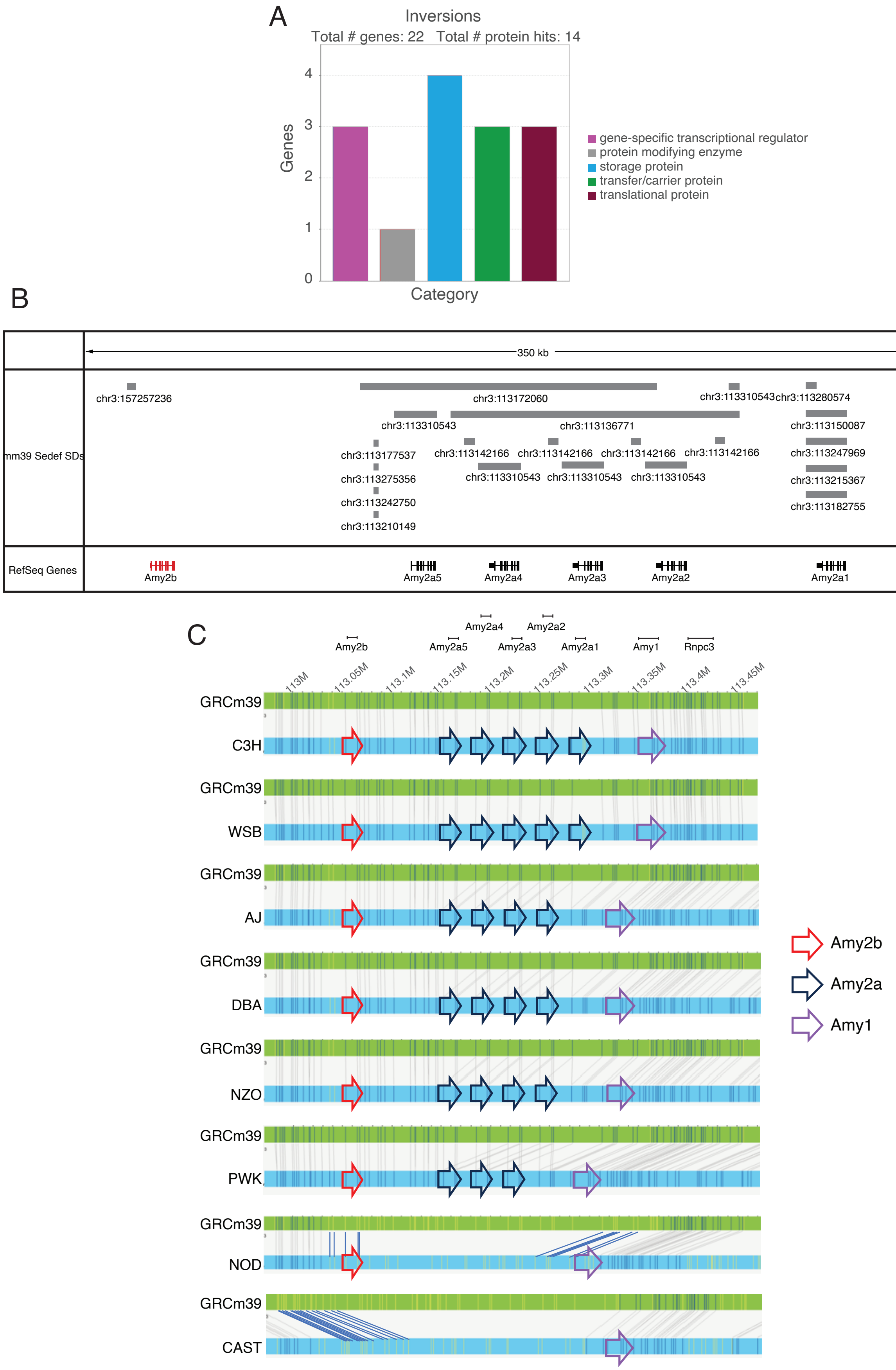


**Supplementary figure 5** (A) PANTHER protein classes of genes that intersect with inversion breakpoints. (B) Amylase locus SD structure in GRCm39. (C) Bionano optical mapping tracks of the amylase locus for all 8 strains. Blue lines connecting GRCm39 track to the respective strain were manually curated.
